# A critical role of a eubiotic microbiota in gating proper immunocompetence in *Arabidopsis*

**DOI:** 10.1101/2023.03.02.527037

**Authors:** Bradley C. Paasch, Reza Sohrabi, James M. Kremer, Kinya Nomura, Yu Ti Cheng, Jennifer Martz, Brian Kvitko, James M. Tiedje, Sheng Yang He

## Abstract

Although many studies have shown that microbes can ectopically stimulate or suppress plant immune responses, the fundamental question of whether the entire preexisting microbiota is indeed required for proper development of plant immune response remains unanswered. Using a recently developed peat-based gnotobiotic plant growth system we found that *Arabidopsis* grown in the absence of a natural microbiota lacked age-dependent maturation of plant immune response and were defective in several aspects of pattern-triggered immunity. Axenic plants exhibited hypersusceptibility to infection by the bacterial pathogen *Pseudomonas syringae* pv. *tomato* DC3000 and the fungal pathogen *Botrytis cinerea*. Microbiota-mediated immunocompetence was suppressed by rich nutrient conditions, indicating a tripartite interaction between the host, microbiota, and abiotic environment. A synthetic microbiota composed of 48 culturable bacterial strains from the leaf endosphere of healthy *Arabidopsis* plants was able to substantially restore immunocompetence similar to plants inoculated with a soil-derived community. In contrast, a 52-member dysbiotic synthetic leaf microbiota overstimulated the immune transcriptome. Together, these results provide evidence for a causal role of a eubiotic microbiota in gating proper immunocompetence and age-dependent immunity in plants.

## Main

The aboveground and belowground parts of land plants host a variety of microorganisms, which collectively constitute the plant microbiota. Microbiota members can reside on or inside plants and appear to be taxonomically conserved at the phylum level^1-8^. The broad conservation of plant microbiota suggests that plants likely have evolved mechanisms to select and maintain the abundance, composition and function of microbiota to achieve homeostasis^9^. A correctly assembled microbiota (i.e., eubiotic microbiota) is likely essential for plant health and survival as recent studies have begun to reveal deleterious effects of genetically induced dysbiotic microbiotas on plant health^10-13^. Although individual or groups of members of the microbiota have been shown to improve nutrient uptake, growth, and resistance to abiotic and biotic stresses^1,2,14-16^, the contribution of a plant’s entire indigenous microbiota to plant functions is not well understood. This is largely due to poorly dissected microbe-microbe and microbe-plant interactions at the community level.

Different members of the plant microbiota can form mutualistic, commensal or pathogenic interactions with plants. To protect against potentially harmful exploitations by microorganisms, plants have evolved cell surface and intracellular immune receptors that recognize evolutionarily conserved microbe-associated molecular patterns (PAMPs) or pathogen-derived effector proteins, resulting in pattern-triggered immunity (PTI) or effector-triggered immunity (ETI), respectively. While ETI appears to be specific for pathogens, PTI represents a basal line of plant defense against both pathogenic and nonpathogenic microbes and is required for maintaining a eubiotic phyllosphere microbiota in *Arabidopsis* to prevent dysbiosis^10,11^. PTI signaling is initiated upon perception of PAMPs by plasma membrane-localized pattern recognition receptors (PRRs)^17^. For example, a 22-amino-acid epitope derived from bacterial flagellin (flg22) is a well characterized elicitor of PTI and recognized by the PRR FLAGELLIN-SENSITIVE 2 (FLS2)^18^. FLS2 forms a complex with co-receptor BRASSINOSTEROID INSENSITIVE 1-ASSOCIATED RECEPTOR KINASE 1 (BAK1)^19^. Phosphorelays between FLS2, BAK1, BOTRYTIS-INDUCED KINASE 1 (BIK1) and a MAPK cascade initiate downstream PTI signaling events, including the production of reactive oxygen species (ROS), calcium fluxes, expression of a large suite of defense-related genes, cell wall remodeling and stomatal closure^20-25^. Activation of PTI prior to an infection can also result in enhanced pathogen resistance^26,27^.

Age-related resistance (ARR) is a widely observed phenomenon in plants in which young plants exhibit greater disease susceptibility compared to older plants^28,29^. This is observed across many flowering plants against a variety of pathogens^30^. In *Arabidopsis*, for instance, the basal susceptibility of young plants to the foliar bacterial pathogen *Pseudomonas syringae* pv. *tomato* (*Pst*) DC3000 is greater compared to older plants^31^. One hypothesis to explain ARR involves the growth-defense tradeoff concept: in order to balance resource allocations during vigorous vegetative growth early in life, young plants prioritize growth over defense^32,33^. Indeed, there is evidence of direct molecular connections between plant growth and immunity^34-36^ including common dual-function signaling components as in the case of PTI and brassinosteroid-dependent plant growth^37^. However, it is unclear whether molecular connections such as these are a sole basis for ARR in plants. In the animal kingdom, development of gnotobiotic animals such as germ-free mice led researchers to discover an important contribution of endogenous microbiota in postnatal maturation of innate immune responses in newborn animals^38,39^. This raises the possibility that plant microbiota may also contribute to the maturation of plant immunity. However, it remains an open question whether age-dependent immunity is entirely intrinsic to plant development or whether maturation of PTI is, in part, the result of colonization of a microbiota. Furthermore, in animals, the presence of dysbiotic microbial communities can be linked to exaggerated immune responses, which have debilitating clinical consequences^40^. Genetically induced and naturally occurring dysbiotic microbial communities have recently been described in plants ^10,41,42^, however it is not clear if dysbiotic microbiota in plants are associated with overactive immune responses. Addressing these basic microbiome questions requires the development of proper gnotobiotic plant growth systems and establishment of well characterized normal (eubiotic) and dysbiotic microbial communities.

In a recent study, we reported two peat-based gnotobiotic plant growth systems, FlowPot and GnotoPot^43^, and two synthetic bacterial communities, a eubiotic community from healthy *Arabidopsis* leaves and a dysbiotic community from leaves of the *Arabidopsis min7 fls2 efr cerk1* (*mfec*) quadruple mutant, which lacks the ability to maintain a eubiotic endophytic bacterial community^10^. Here, we employed these tools to address the questions regarding the role of the endogenous microbiome in the development of ARR and a possible role of eubiosis in gating proper plant basal immunity.

## Results

### Age-dependent PTI in conventionally grown plants

We began this project by characterizing possible maturation of PTI over time in conventionally grown *Arabidopsis* plants. For this purpose, we performed the classical flg22 protection assays using 2.5-week-old and 3.5-week-old *Arabidopsis* plants, which were conventionally grown in a potting soil substrate in air-circulating growth chambers. In 2.5-week-old plants, we observed a modest level of flg22-mediated resistance to virulent *Pst* DC3000 in flg22-treated plants, compared to mock-treated plants. Older, 3.5-week-old plants, however, exhibited a significantly enhanced level of flg22-triggered resistance, compared to 2.5-week-old plants (Fig. 1a). This result demonstrated age-dependent development of PTI in soil-grown *Arabidopsis* plants, which is consistent with a recent study showing that FLS2-dependent immunity increased in the first 6 days of young seedling growth in agar medium without microbes^44^.

**Figure 1.**
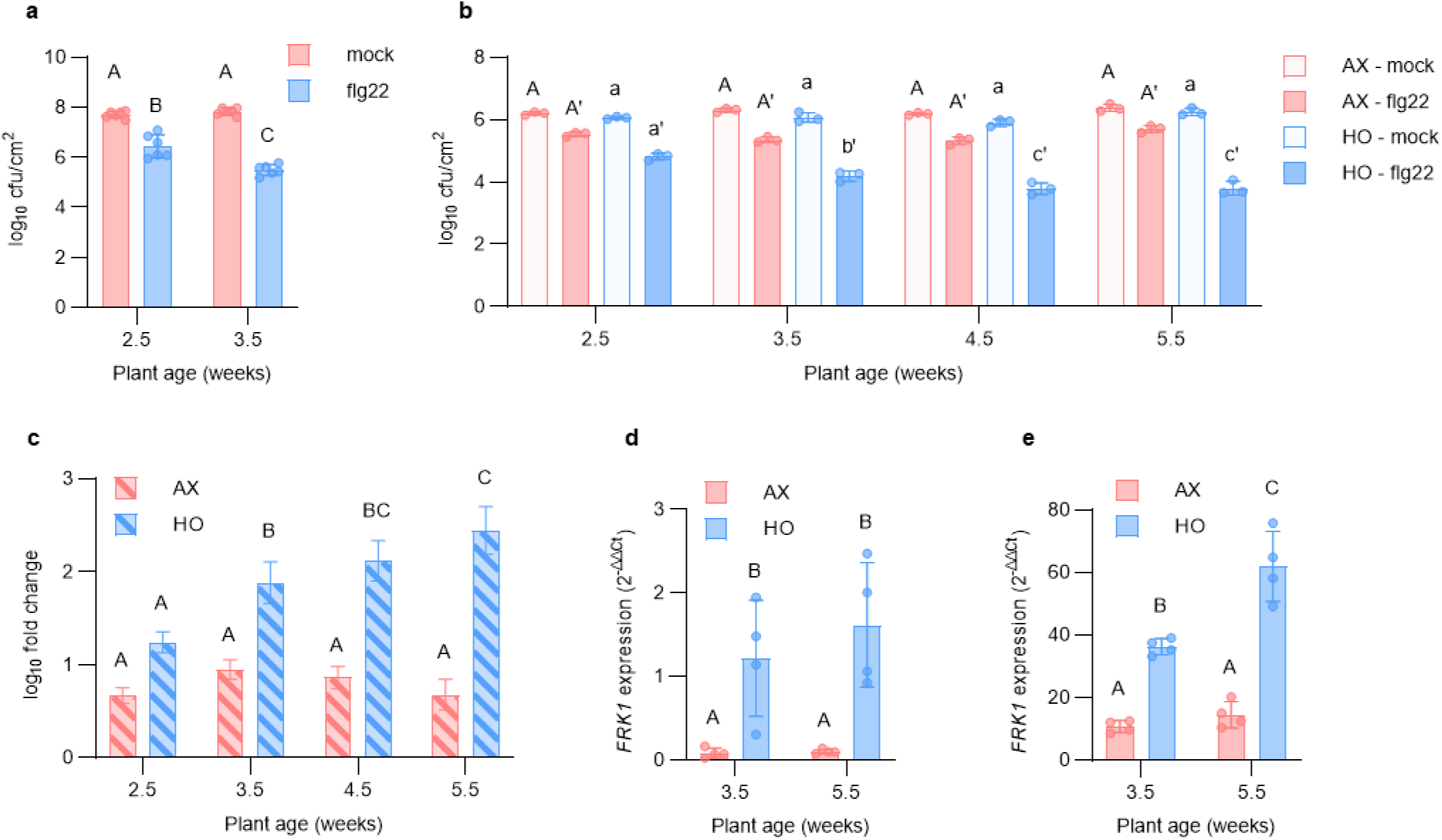
Age-dependent flg22-triggered immunity in *Arabidopsis*. **a**, flg22 protection assay showing enhanced resistance against *Pst* DC3000 triggered by pretreatment with 500 nM flg22 in 2.5-week-old and 3.5-week-old plants. Each column represents bacterial titer 24 hours after noculation as log transformed colony forming units (cfu)/cm^2^ and is the mean of six plants. Error bars indicate SD. Different letters represent a significant difference (*p* < 0.05, two-way ANOVA with Tukey’s HSD post-hoc test). **b**, Age-dependent flg22 protection. Axenic (AX) or holoxenic (HO) plants were treated 24 hours prior to inoculation with *Pst* DC3000 with either a water (mock) or 100 nM flg22 solution. Each column represents bacterial titer 24 hours after inoculation as log transformed cfu/cm^2^ and is the mean of 3 plants. Error bars indicate SD. Different letters represent a significant difference (*p* < 0.05, two-way ANOVA with Tukey’s HSD post-hoc test). **c**, Relative protection displayed as fold change in bacterial cell counts between flg22- and mock-treated samples. Error bars represent SD. Different letters represent a significant different (*p* < 0.05, two-way ANOVA with Tukey’s Honest Significant Difference (HSD) post-hoc test). **d**, Basal and **e**, flg22-induced age-dependent *FRK1* gene expression in 3.5-week-old and 5.5-week-old AX and HO plants. Total RNA was extracted 4 h after treatment with a mock solution lacking flg22 for basal expression or 100 nM flg22 for flg22-induced expression. Expression levels displayed as relative to mock treated 3.5-week-old HO plants for both panels. *PP2AA3* was used for normalization. Results represent the mean values ± SD of four plants. Different letters represent a significant difference (*p* < 0.05, two-way ANOVA with Tukey’s HSD post-hoc test). Each experiment was repeated three independent times with similar results.

### Age-dependent PTI maturation requires microbiota colonization

Traditionally, age-related resistance has been attributed to developmental transition processes^45^. We examined an additional hypothesis that the endogenous microbiome might be involved in age-dependent PTI in plants. For this purpose, we investigated the temporal maturation of flg22-mediated resistance in peat-based gnotobiotic plant growth systems. Holoxenic (HO) plants colonized with a natural, soil-derived microbial community (‘MSU’, which was harvested from agricultural soil located at Michigan State University, East Lansing, Michigan; see Methods) and corresponding axenic (AX) plants, which were mock-inoculated with autoclaved ‘microbial community derived from the same ‘MSU’ soil, were used. As shown in Fig. 1b, HO plants exhibited progressively more robust flg22-mediated resistance against *Pst* DC3000 over time, which is consistent with age-dependent PTI observed in plants grown conventionally in potting soil (Fig. 1a). In contrast, AX plants mock-inoculated with the autoclaved microbial community were greatly reduced in age-dependent flg22-mediated resistance phenotype (Fig. 1c). *Arabidopsis* mutant *bak1-5 bkk1-1 cerk1* (*bbc)*^46^; which is defective in PTI signaling downstream of multiple PRRs, including the flg22 receptor FLS2, did not show flg22-mediated resistance in HO plants at any age (Supplementary Fig. 1), suggesting that microbiota-mediated age-dependent resistance requires canonical PTI signaling co-receptors.

Next, we quantified induction of the PTI marker gene *FLG22-INDUCED RECEPTOR-LIKE KINASE 1* (*FRK1*) to further characterize age-dependent activation of PTI in HO plants and apparent lack thereof in AX plants. While basal expression of *FRK1* was similar for both 3.5- and 5.5-week-old HO plants (Fig. 1d), flg22 induced a higher level of *FRK1* expression in old HO plants than in younger HO plants (Fig. 1e). Interestingly, basal expression of *FRK1* was lower in AX plants compared to either young or old HO plants (Fig. 1d) and, notably, no significant age-dependent increase in flg22-induced *FRK1* expression was observed in AX plants (Fig. 1e). Thus, the reduced age-dependent maturation of PTI in AX is correlated with a lack of robust increase in age-dependent expression of *FRK1* gene.

### Axenic plants lack normal expression of immune-associated genes

To capture genome-wide gene expression in AX and HO plants beyond the *FRK1* marker gene, we conducted transcriptome analysis of AX and HO *Arabidopsis* plants grown in the peat gnotobiotic system. To reduce the possibility of community-specific bias due to use of a single microbiota, microbial communities harvested from two distinct soils were used: ‘MSU’, which was harvested a Alfisol soil type, and ‘Malaka’, which was harvested from undisturbed grassland soil in Malaka Township, Iowa (see Methods) and is a Mollisol soil. Principal component analysis (PCA) of RNA-seq gene expression data revealed distinct expression patterns between HO plants and AX plants (PC1, 26% variance; Fig. 2a). Using a |log_2_FC| ≥ 1 and false discovery rate (FDR) < 0.05 cutoff, we identified a total of 435 differentially expressed genes (DEGs) between HO and AX plants across both microbiota inputs: 352 were depleted in AX plants and 83 were enriched in AX plants (Fig. 2b,c and Supplementary Table 1). Of the 352 DEGs depleted in AX plants, 138 were depleted irrespective of the microbiota input source (i.e., enriched in both HO plants colonized by the ‘MSU’ community and HO plants colonized by the ‘Malaka’ community; Fig. 2d). Gene ontology (GO) term enrichment analysis of these 138 ‘core’ AX-depleted genes revealed an overrepresentation of terms involved in plant immunity (Fig. 2e and Supplementary Table 2). The genes enriched in AX plants did not display any significant GO term enrichment. Closer examination of depleted DEGs in AX plants revealed numerous genes involved in PTI, SA-mediated defense, and defense-associated metabolite biosynthesis (Fig. 2c and Supplementary Table 1). These genes included *FRK1*; several leucine-rich repeat protein kinases such as *IMPAIRED OOMYCETE SUSCEPTIBILITY 1* (*IOS1*), *AT1G51890*, *AT1G51790, AT1G51860* and *AT5G59680*; systemic immunity-associated genes *AZELAIC ACID INDUCED 1* (*AZI1*) and *AZI3*; *PATHOGENESIS RELATED 2* (*PR2*) and *PR4;* glucosinolate biosynthesis genes such as *FAD-LINKED OXIDOREDUCTASE* (*FOX1*) and the cytochrome P450 monooxygenases *CYP71A12 and CYP71B15*; and defense-associated transcription factors *MYB15* and *WRKY 30* (Fig 2c and Supplementary Table 1). Thus, consistent with the targeted *FRK1* gene expression analysis shown in Fig. 1d, results from the transcriptome analysis using two independent soil-derived microbiotas pointed to a broadly depleted PTI/SA defense gene expression in AX plants compared to HO plants, which collectively contribute to induced and basal innate immunity.

**Figure 2.**
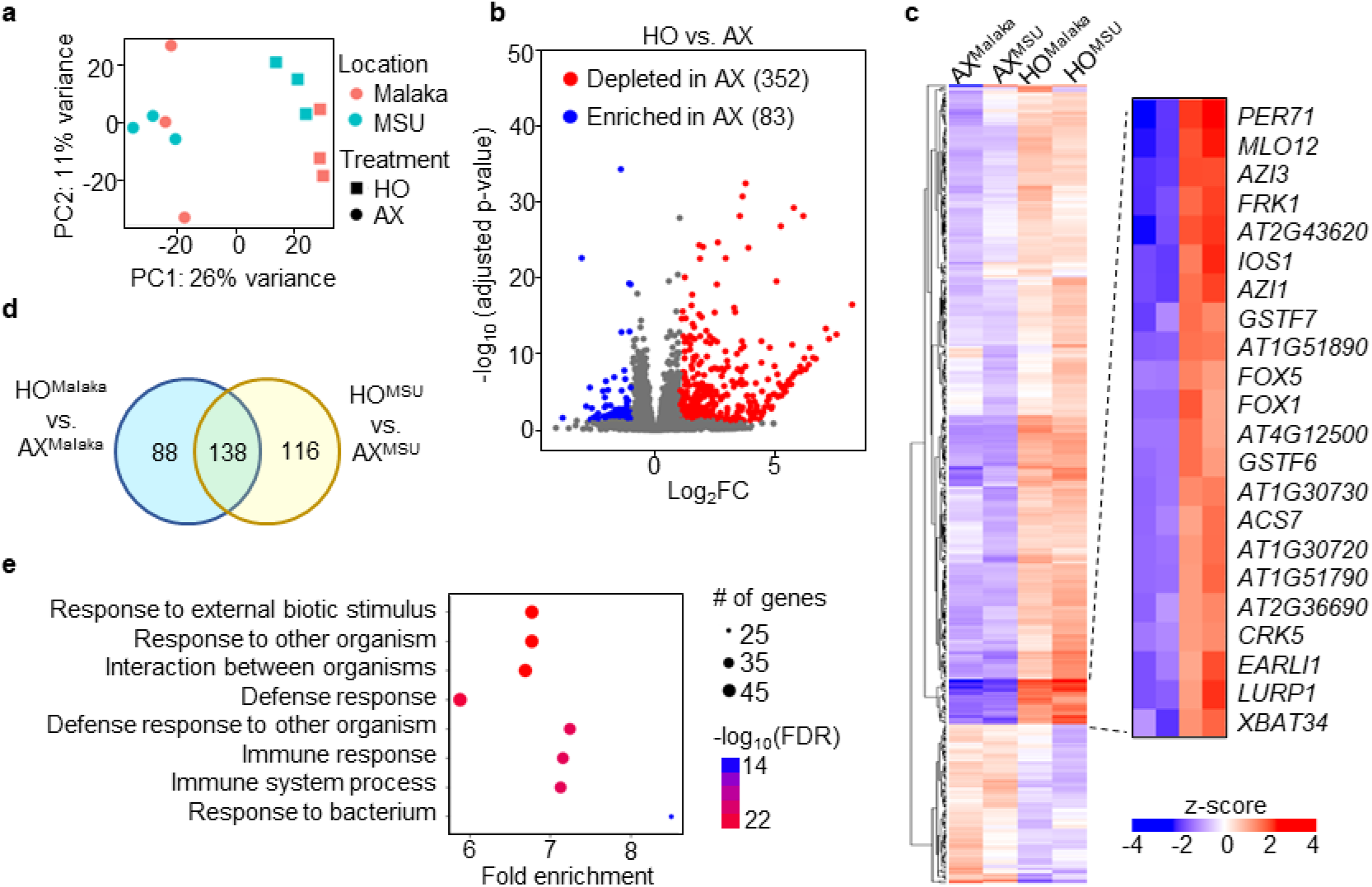
Axenic *Arabidopsis* plants are depleted in the basal expression of defense-related transcripts. **a**, Principal component (PCA) analysis of genes expressed under AX and HO conditions using microbial communities from two different locations/soil types (‘MSU’: MI, Alfisol soil type and ‘Malaka’: IA, Mollisol soil type; 8 plants from one microbox pooled per biological rep, n = 3). **b**, Volcano plot of differentially expressed genes (DEGs). Colored regions represent significant differential expression with |log_2_FC| > 1 and FDR < 0.05 (Benjamini-Hochberg corrected Wald Test) with the number of genes corresponding to each group indicated in parentheses. **c**, Heat map of DEGs generated using hierarchical clustering with euclidean distance and complete linkage. Label superscript indicates community used for inoculation of HO plants or mock inoculation of AX plants. A subset of the differentially regulated genes in HO and AX is hown on right. **d**, Venn diagram of upregulated DEGs showed 138 common genes in response to HO^MSU^ and HO^Malaka^ treatments. **e**, Gene ontology (GO) term enrichment (GO:BP biological process) analysis on ‘core’ depleted DEGs in AX plants. Top enriched GO terms displayed, ranked by significance (FDR adjusted hypergeometric test).

### Axenic *Arabidopsis* is underdeveloped in PTI

In addition to depleted immune gene expression, we found AX plants exhibited significantly lower levels of other PTI-associated immune responses compared to HO plants. For example, 6-week-old AX plants exhibited significantly reduced flg22-, elf18-, and Pep1-induced ROS production compared to HO plants both in the magnitude of maximum ROS production (peak amplitude) and in the time to reach the maximum (Fig. 3a and Supplementary Fig. 2). AX plants also exhibited significantly reduced PAMP/DAMP-induced *FRK1* gene expression compared to HO plants (Fig. 3b). Western blot analysis revealed that, despite possessing similar levels of total MPK3 and MPK6 (Fig. 3c,d), less MPK was phosphorylated in AX plants after the activation of PTI by treatment with flg22 (Fig. 3e). Although RT-qPCR analysis consistently showed that both basal and flg22-induced expression of the *FLS2* receptor gene is significantly reduced in AX plant leaf tissue compared to HO plant leaf tissue (Fig. 3f), total FLS2 protein abundance was variable and only occasionally reduced in AX plant leaves (Supplementary Fig. 3). In contrast, the co-receptor BAK1 protein was consistently found in lower relative abundance in AX plants compared to HO plants (Fig. 3g). Additionally, quantification of the defense hormone salicylic acid (SA), which is downstream of PTI signaling, revealed that AX plants possess lower basal levels of SA compared to HO plants (Fig. 3h,i). Finally, AX plants were hypersensitive to infection by the virulent foliar hemibiotrophic bacterial pathogen *Pst* DC3000 and the necrotrophic fungal pathogen *Botrytis cinerea*, compared to HO plants (Fig. 3j,k). Together, these studies demonstrate multiple compromised PTI immune phenotypes in axenic plants.

**Figure 3.**
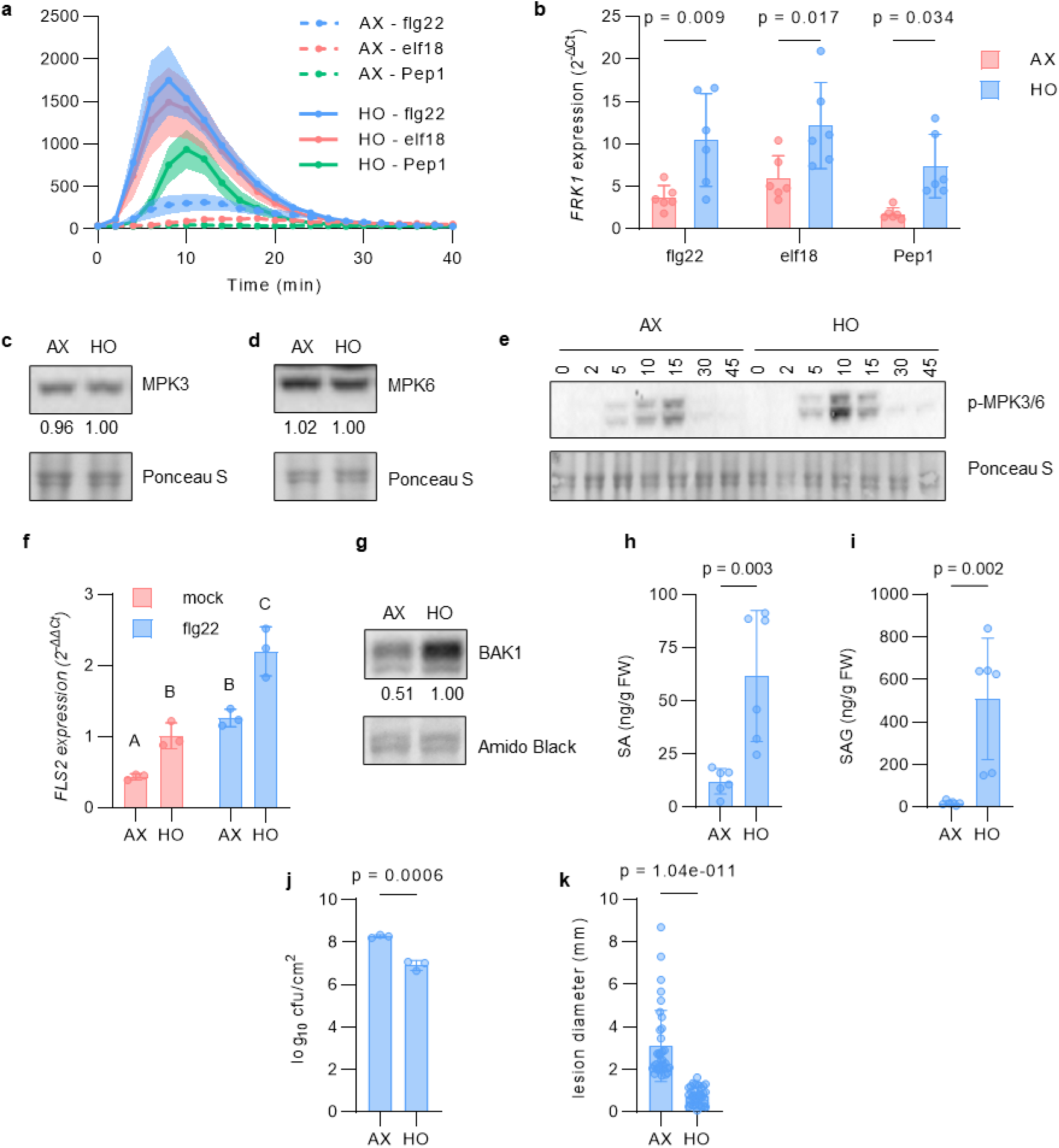
Axenic *Arabidopsis* plants exhibit defects in PTI compared to colonized plants. **a**, ROS burst dynamics induced by 250 nM flg22, elf18, and Pep1 in AX and HO plants in GnotoPots. Results represent the mean of eight plants ± SEM. **b**, *FRK1* gene expression in AX and HO plants induced by 250 nM flg22, elf18, and Pep1. Total RNA was extracted from leaf disks 1.5 h after treatment. *PP2AA3* was used for normalization. Bars represent the mean value of eight plants ± SD. (flg22: *p* = 0.009, elf18: *p* = 0.017, Pep1: *p* = 0.034; Two-way ANOVA with Šidák multiple comparison test). **c**,**d**, Representative blots of total MPK3 (c) or MPK6 (d) proteins in 4.5 week old AX and HO plants. Protein was detected with MPK3 or MPK6-specific antibodies. **e**, Representative blot of phosphorylated MPK3/6 proteins detected using an α-p44/42-ERK antibody upon treatment with 100 nM flg22. Samples were taken at the indicated times after treatment. For panels (c,d), numbers below blots indicate band intensity relative to that of Ponceau S, normalized to HO = 1.00. **f**, Basal and flg22-induced expression of *FLS2* gene in AX and HO plant leaf tissue. Total RNA was extracted 1 h after treatment with 100 nM flg22 or mock solution. PP2AA3 was used for normalization. Bars represent the mean values ± SD of three biological samples, each consisting of three plants. Different letters represent a significant difference (*p* < 0.05, unpaired *t*-tests with Welch’s correction). **g**, Total BAK1 protein detected in leaf lysates of AX and HO plants. Numbers below blot indicates band intensity relative to that of Amido Black, normalized to HO = 1.00. **h**,**i**, Total levels of salicylic acid (SA) (h) and glucosylated SA (i) in AX and HO plants. Each bar represents the mean values ± SD of eight biological samples, each consisting of at least three plants. (SA: *p* = 0.003, SAG: *p* = 0.002; Student’s t-test). **j**, *Pst* DC3000 populations in AX and HO plants. Each column represents bacterial titer 3 days after inoculation as log transformed cfu/cm^2^ and is the mean of three plants. Error bars indicate SD (*p* = 0.0006, Student’s *t*-test). **k**, Size of lesions formed in AX and HO plants by *Botrytis cinerea* (*Bc*). Each column represents lesion diameter 5 days after inoculation and is the mean of 36 lesions on six plants. Error bars indicate SD (*p* = 1.04 × 10^-11^, Student’s *t*-test). Each experiment was repeated three independent times with similar results. See Source Data Figures 1-4 for image cropping.

### A eubiotic leaf synthetic community (SynCom) substantially confers immunocompetence

We recently assembled a 48-member eubiotic SynCom (SynCom^Col-0^) composed of endophytic bacteria from leaves of healthy *Arabidopsis* Col-0 plants^10^. To determine to what extent an eubiotic SynCom derived from the leaf endosphere could restore immunocompetence to AX plants, we compared the PTI phenotypes of Col-0 plants grown with and without SynCom (SynCom^Col-0^ vs. MgCl_2_). Col-0 plants grown with the ‘MSU’ soil-derived microbiota were used as control. We observed robust flg22-induced production of ROS in HO plants inoculated with the ‘MSU’ soil-derived microbiota and SynCom^Col-0^-inoculated plants (Fig. 4a and Supplementary Fig. 4a). We next quantified flg22-induced *FRK1* gene expression and observed that plants colonized by SynCom^Col-0^ were restored in basal and flg22-induced *FRK1* expression (Fig. 4b,c), which was again similar to what was observed for HO plants (Fig. 1d,e). Additionally, plants colonized by SynCom^Col-0^ had an increased level of BAK1 protein (Supplementary Fig. 4b) and were more resistant to *Pst* DC3000 infection (Fig. 4d) compared to AX plants mock-inoculated with the same volume of 10 mM MgCl_2_. Taken together, these results suggest that a leaf endosphere-derived bacterial SynCom can substantially restore immune competence to AX plants similar to a natural soil-derived microbiota.

**Figure 4.**
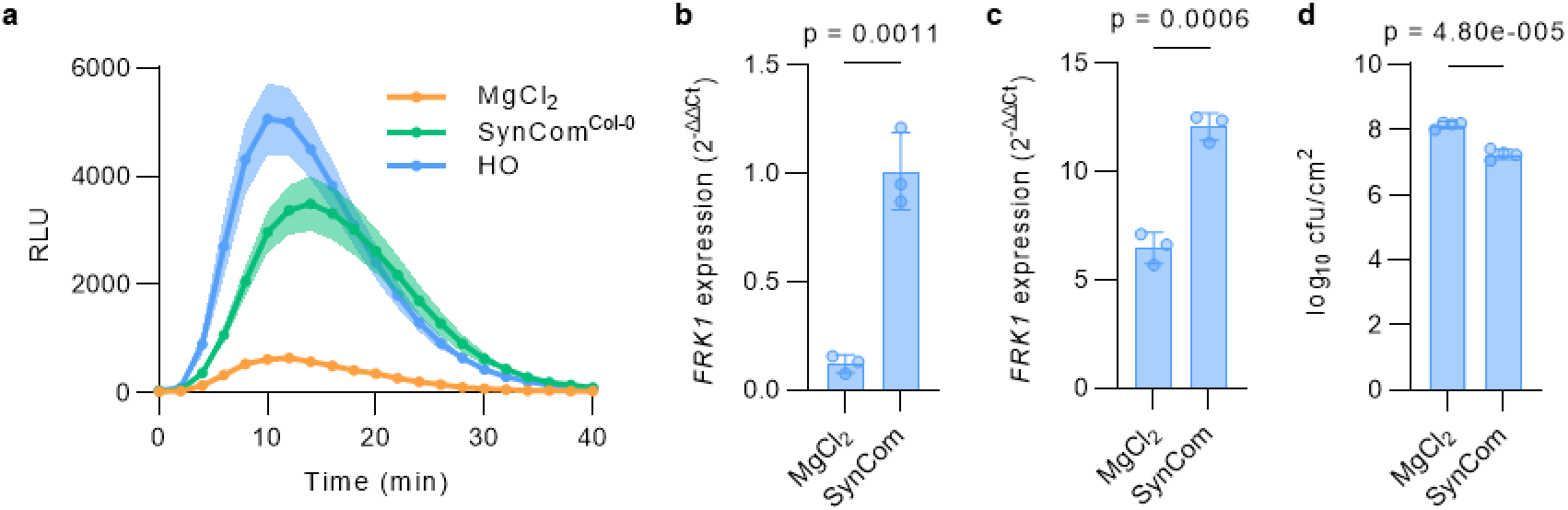
Natural microbiota and SynCom^Col-0^ restore immunocompetence. **a**, ROS burst dynamics induced by 100 nM flg22 in axenic plants mock-inoculated with 10 mM MgCl_2_ and plants colonized by HO or SynCom^Col-0^. Results represent the mean of 12 plants ± SEM. **b**,**c**, Basal (b) and flg22-induced (c) *FRK1* expression in axenic MgCl_2_ mock-inoculated plants and plants inoculated with SynCom^Col-0^. Total RNA was extracted 3 h after treatment with a mock solution lacking flg22 (b) or 100 nM flg22 (c). Results relative to basal expression in SynCom^Col-0^-inoculated plants. *PP2AA3* was used for normalization. Bars represent the mean values ± SD of three plants. Different letters represent a significant difference. Asterisks represents a significant difference (basal expression: *p* = 0.0011, flg22-induced: *p* = 0.0006, Student’s *t*-test). **d**, *Pst* DC3000 populations in axenic plants mock-inoculated with 10 mM MgCl_2_ and SynCom^Col-0^-inoculated plants. Each column represents bacterial titer 3 days after inoculation as log transformed cfu/cm^2^ and is the mean of four plants. Error bars indicate SD (*p* = 4.80 × 10^-5^, Student’s *t*-test).

To evaluate possible redundancy of SynCom^Col-0^ members in contribution to immune competence we assembled a simplified version of SynCom^Col-0^ with only 19 strains (SynCom^Col-mix19^) to cover the taxonomic diversity at the genus level (Supplementary Table 3). We found SynCom^Col-mix19^ could effectively restore ROS production in response to flg22 (Supplementary Fig. 5). Furthermore, among strains with high representation in SynCom^Col-0^, we randomly chose a mix of three strains which are present in SynCom^Col-mix19^ and found that these three strains (SynCom^Col-mix3^, representing *Achromobacter*, *Comamonas* and *Stenotrophomonas* genera) also partially restored ROS production in response to flg22 (Supplementary Fig. 5). This suggest that there might be significant redundancy among strains of SynCom^Col-0^ that can endow immunocompetence.

### Impact of abiotic conditions on microbiota-mediated immune competence

During the development and optimization of the peat-based gnotobiotic system, we noticed a correlation between levels of microbiota-mediated restoration of immune competency and concentrations of Linsmaier and Skoog^47^ (LS) nutrient media (which contains mineral slats as well as some organic compounds such as myo-inositol and MES buffer; see Methods for nutrient content) added during preparation of the gnotobiotic system. To systematically determine the effects of nutrients on microbiota-mediated immunocompetence we measured flg22-induced production of ROS in AX and HO plants along a nutrient concentration gradient. Using the same volume of liquid, GnotoPots were prepared with full strength (1x) LS, half strength (0.5x) LS and one tenth strength (0.1x) LS. We observed a significant impact of nutrients on flg22-mediated ROS production in HO plants. Decreasing nutrient strength significantly increased ROS burst magnitude and shortened time to reach the maximum ROS production (Fig.4e) in HO plants. At intermediate nutrient levels (0.5x LS), ROS burst magnitude was moderately increased and time to reach the maximum was reduced compared to higher (1x LS) nutrient concentrations, however the total ROS produced was not significantly different (Fig. 5a and Supplementary Fig. 6a). At low nutrient levels (0.1x LS), ROS burst magnitude was increased, time to maximum shortened, and total ROS increased (Fig. 5a and Supplementary Fig. 6a). Nutrient concentration did not have an effect on the timing of ROS production in AX plants, and only a marginal impact on total ROS production was observed. We next examined the effects of individual components of LS medium, including nitrogen, phosphorus, and iron as well as carbon containing compounds myo-inositol and MES, on microbiota-mediated immunocompetence by supplementing 0.5x LS with each nutrient/compound at the concentration present in 1x LS. Only 1x nitrogen (supplied as NH_4_NO_3_ and KNO_3_ salts) suppressed flg22-induced ROS production and *FRK1* gene expression in HO plants to levels similar to 1x LS (Supplementary Fig 6b-d), indicating high N content plays a major role in suppressing microbiota-mediated immunocompetence.

**Figure 5.**
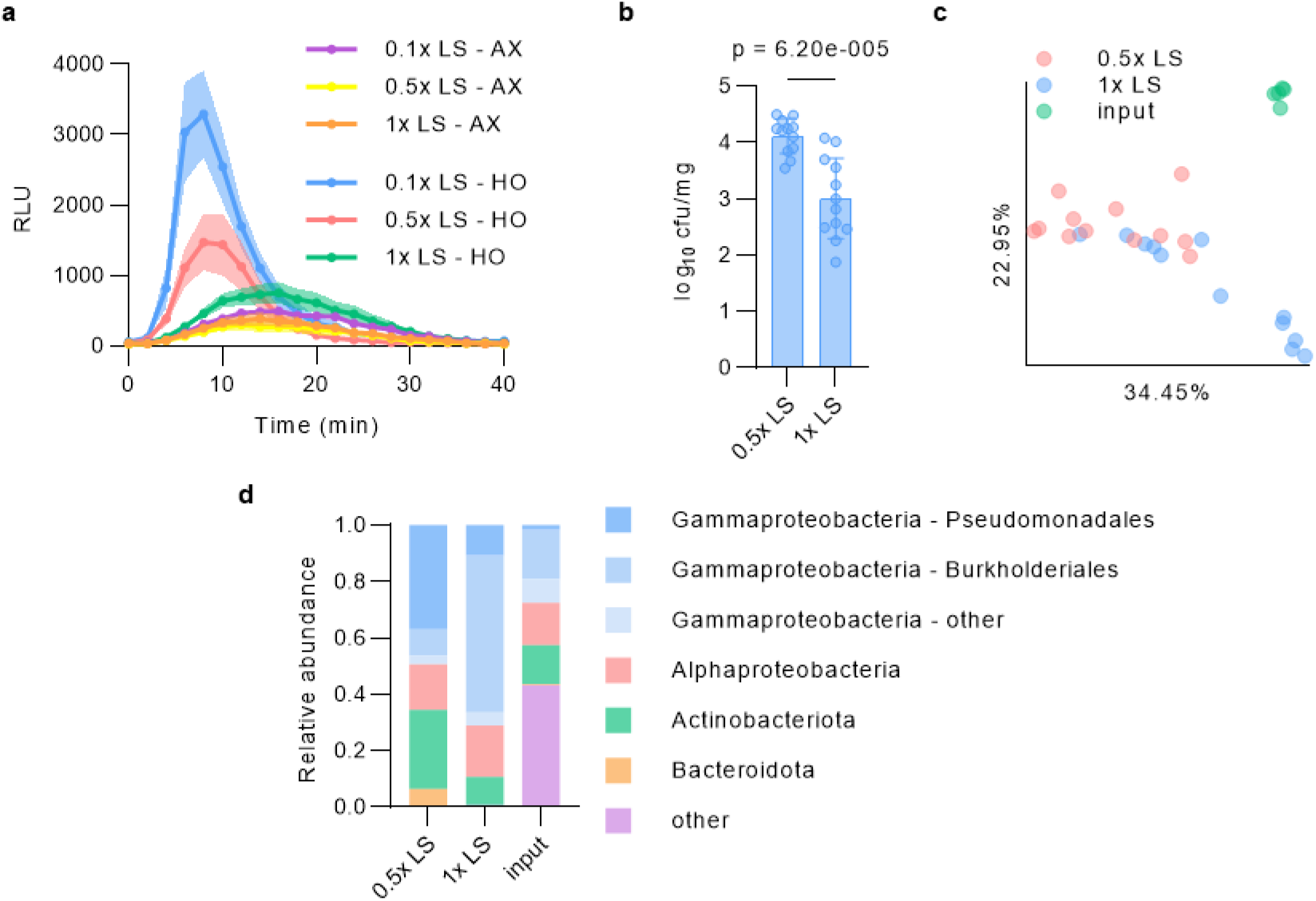
Microbiota-mediated immunocompetence is nutrient-dependent. **a**, ROS burst dynamics induced by 100 nM flg22 in AX and HO plants grown in GnotoPots supplied with 0.1x, 0.5x, or 1x LS nutrient solution concentrations. Results represent the mean of six plants ± SEM. **b**, Absolute abundance of phyllosphere bacterial populations associated with HO plants grown in GnotoPots supplied with either 0.5x or 1x LS nutrient solution. Each column represents bacterial titer as log transformed cfu/cm^2^ and is the mean of 12 plants. Error bars indicate SD (*p* = 6.20 × 10^−#1^, Student’s *t*-test). For panels (a-g), experiments were repeated a minimum of two independent times with similar results. **c**, PCoA of weighted UniFrac distances obtained from 16S rRNA gene sequence profiles of soil-derived input microbiota (MSU) and phyllosphere microbiota of HO plants after 6 weeks of growth in GnotoPots supplied with either 0.5x or 1x LS nutrient solution. (0.5x LS vs. 1x LS: *q* = 0.002, 0.5x LS vs. input: *q* = 0.002, 1x LS vs. input: *q* = 0.002; pairwise PERMANOVA with 999 permutations). **d**, Relative abundance of bacterial populations at the phylum level. Members of Proteobacteria phylum are separated into class and members of Gammaproteobacteria class are further separated into order. *n* = 5 (input), *n* = 12 (0.5x LS), *n* = 12 (1x LS).

To determine whether microbial colonization is affected by nutrient level, we determined the absolute and relative abundance of phyllosphere bacterial microbiota using enumeration of culturable bacteria and 16S rRNA gene amplicon sequencing in plants supplemented with 0.5x LS or 1x LS. Plants grown with 1x LS harbored approximately 10-fold lower total phyllosphere bacteria microbiota levels compared to plants grown with 0.5x LS (Fig. 5b). Principal coordinates analysis (PCoA) on weighted UniFrac distances indicated a significant compositional difference between phyllosphere bacterial communities associated with plants grown under the two nutrient levels (Fig. 5c). Actinobacteriota, Bacteroidota and Gammaproteobactera (belonging to the order Pseudomonadales) were more abundantly observed in the phyllosphere of plants grown with 0.5x LS, whereas their relative abundance was greatly reduced in plants grown with 1x LS. Conversely, Gammaproteobacteria belonging to order Burkholderiales increased relative abundance in plants grown with 1x LS compared to those grown with 0.5x LS (Fig. 5d). Together, these findings illustrated a tripartite interaction between immunity, microbiota, and environment during microbiota-mediated maturation of flg22-triggered immunity.

### Dysbiotic microbiota overstimulates immune gene expression

Several recent reports have begun to show an important contribution of plant immunity, including PTI and vesicular trafficking pathways, to maintaining microbiota homeostasis in Arabidopsis leaves^1,10,11^. In particular, we were able to establish two parallel leaf endosphere-derived bacterial SynComs: 48-member SynCom^Col-0^ derived from healthy Col-0 leaves and 52-member SynCom^mfec^ derived from dysbiotic *mfec* mutant leaves^10^. To investigate the impact of a normal (eubiotic) microbiota vs. a dysbiotic microbiota on plant immunity, we examined the expression of several immunity-associated marker genes (*FRK1*, *PR1* and *CYP71A12*) in plants colonized with SynCom^mfec^ or SynCom^Col-0^ in comparison to AX plants in plate-based gnotobiotic system. We found a gradient of expression of these genes, with the highest expression observed in Col-0 plants colonized by SynCom^mfec^, an intermediate level in SynCom^Col-0^-colonized plants and the lowest level in AX plants (Fig. 6a-c).

**Figure 6.**
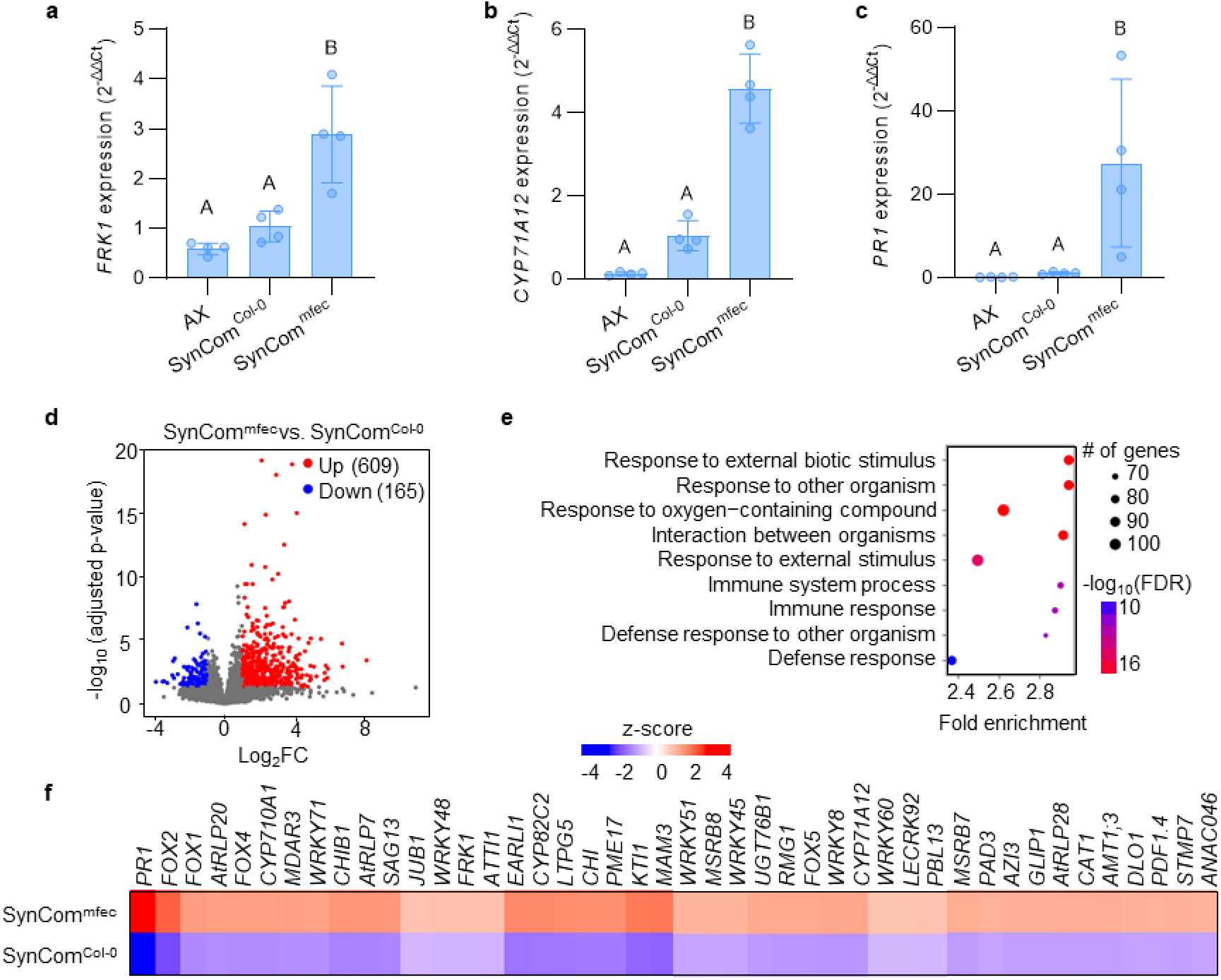
Dysbiotic microbiota overstimulates immune gene expression. **a-c**, Basal expression of defense-related genes *FRK1* (a), *CYP71A12* (b), *PR1* (c) in AX, SynCom^Col-0^- and SynCom^mfec^-inoculated plants grown in agar plates. *PP2AA3* was used for normalization. Bars represent the mean values ± SD of three biological samples, each consisting of five seedlings. Different letters represent a significant difference (*p* < 0.05, one-way ANOVA with Tukey’s HSD post-hoc test). **d**, Volcano plot of genes differentially expressed in SynCom^Col-0^- and SynCom^mfec^-colonized plants. Colored regions represent significant differential expression with |log_2_FC| > 1 and FDR < 0.05 (Benjamini-Hochberg corrected Wald Test) with the number of genes corresponding to each group indicated in parentheses. **e**, GO term enrichment for up-regulated DEGs in SynCom^mfec^-colonized plants compared to SynCom^Col-0^-colonized plants, ranked by significance (FDR adjusted hypergeometric test). **f**, Heat map for selected genes from hierarchical clustering all DEGs. Gene descriptions are listed in supplementary table 4.

To gain a better understanding of plant transcriptional responses to eubiotic microbiota vs. a dysbiotic microbiota, we performed RNA-seq analysis of Col-0 plants colonized by SynCom^mfec^ and SynCom^Col-0^ grown in parallel in the GnotoPot system. Colonization with SynCom^Col-0^ compared to SynCom^mfec^ resulted in 774 DEGs (|log_2_FC| > 1 and FDR < 0.05) (Fig. 6d and Supplementary Table 4). GO term analysis of the 609 DEGs upregulated upon colonization with SynCom^mfec^ vs. SynCom^Col-0^ showed an over-representation of GO terms associated with biotic stress and immunity (Fig. 6e and Supplementary Table 5). Additionally, several immunity pathways including the systemic acquired resistance, PTI signaling and glucosinolate biosynthetic processes were upregulated. Further analysis showed that several dysbiosis-associated genes were involved in pathogenesis-related processes during biotic stresses which are associated with immunity, cell death and its regulation (Fig. 6f). Collectively, our results showed that dysbiotic SynCom^mfec^ overstimulates immune gene expression compared to eubiotic SynCom^Col-0^.

Next, we examined the capacity of individual SynCom members to potentiate immune stimulation. To facilitate the analysis of immune gene expression involving a large number of microbiota strains (48 SynCom^Col-0^ strains and 52 SynCom^mfec^ strains), we first performed qualitative GUS assays with 12-day-old seedlings of the *CYP71A12_Pro_:GUS* reporter line grown in liquid LS media inoculated with each of the 100 individual SynCom members. We found that the *Stenotrophomonas maltophilia* strains from both SynCom^Col-0^ (4 strains) and SynCom^mfec^ (8 strains) induced *CYP71A12_Pro_:GUS* reporter in leaves. Additionally, 4 other strains that are unique to SynCom^mfec^ including *Stenotrophomonas acidaminiphila* (mfec-41), *Stenotrophomonas sp*.(mfec-48), *Microbacterium Sp* (mfec-31) and *Pseudomonas citronellolis* (mfec-34), showed *CYP71A12_Pro_:GUS* reporter activity in seedling leaves (Supplementary Fig. 7). Thus, SynCom^mfec^ has higher number and more diverse strains that have the ability to induce *CYP71A12* promoter activity in leaves. We then performed an independent RT-qPCR based analysis of *CYP71A12* gene expression in leaves of 5 week-old, soil-grown Arabidopsis Col-0 plants, revealing a pattern of *CYP71A12* gene expression similar to *CYP71A12_Pro_:GUS* reporter assay, despite very different plant growth conditions in these two independent experiments (Supplementary Table 6). Notably, most of *CYP71A12*-induced SynCom members were previously shown to cause dysbiotic symptoms^10^.

## Discussion

Here, we show that *Arabidopsis* plants grown without exposure to a microbiota are greatly compromised in age-dependent immunity that occurs in plants colonized naturally by microbiota. Axenically-grown plants exhibit significant defects in PTI and are hypersusceptible to infection by the bacterial pathogen *Pst* DC3000 and the fungal pathogen *B. cinerea*. We also show that immunocompetence can be restored by natural soil-derived microbiota as well as a 48-member eubiotic bacterial synthetic community (SynCom^Col-0^) derived from leaf endophytic bacteria. In contrast, a 52-member dysbiotic synthetic community derived from leaf endophytic bacteria overstimulates immune gene expression. Finally, our results show that the immune-modulation function of microbiota can be influenced by environmental conditions. Together, these results have significant implications in the formulation of a framework for explaining age-dependent immunity, microbiota-immunity interplay and ‘immunity-microbiome-environment’ tritrophic interactions in plants.

With respect to age-dependent immunity, a previous study characterized the ontogeny of flg22-triggered immunity in very young *Arabidopsis* seedlings (within six days after germination) in axenic nutrient agar plates^44,48^, providing insight into the developmentally controlled maturation of immune responses immediately after germination. Results presented here, however, show that flg22-triggered immunity exhibits an age-dependent maturation period that extends through at least the first 2-3 weeks of vegetative growth and that full-scale age-dependent immune maturation requires exposure to microbiota. As demonstrated here, microbiota-colonized HO plants in peat-based gnotobiotic systems developed age-dependent PTI over time, mirroring plants grown conventionally in potting soil. In contrast, development of age-dependent PTI was greatly reduced in AX plants. The microbiota-mediated age-dependent maturation bears striking conceptual parallels to that observed in germ-free mice in which an important contribution of endogenous microbiota in postnatal maturation of mammalian innate immunity is well recognized^38,39^. While ARR has typically been proposed to be caused by developmental processes that antagonize immune responses^45^, results presented here revealed that microbiota-assisted immune maturation is a previously unrecognized contributor that plays an important role in age-dependent immune maturation in plants.

It should be pointed out that the discovery of a causal role of microbiota in age-dependent immune maturation required the use of a gnotobiotic system capable of growing plants with or without a natural or synthetic community of microbes. Because agar plates, a commonly used gnotobiotic system, is not ideal for natural colonization of plants by a complex microbial community due to artificial overgrowth of some microbes, this has been achieved in this study by using FlowPot and GnotoPot gnotobiotic systems with a peat-based substrate, which partially simulates the natural soil substrate. We used FlowPots and GnotoPots interchangeably and some initial experiments were repeated using both systems with similar results. For most subsequent experiments we used GnotoPots, because they allowed plants to grow for a longer duration compared to FlowPots^43^. An important realization during this study is that peat-based plant gnotobiotic systems can be fine-tuned to simulate a range of various abiotic conditions, such as nutrients. This was useful for our study because many of the microbiota functions in nature seem to be context dependent. For example, high nitrogen fertilizer regimes have been shown to increase susceptibility of plants grown in a non-sterile hydroponic system^49^. However, it was not known if the effect of high nitrogen nutrients is mediated in part by microbiota. In this study, fine-tuning the nutrient conditions of GnotoPots enabled us to discover that nutrient-mediated immune suppression was most obvious in the presence of microbiota and that high nitrogen has a prominent effect on microbiota level and composition, suggesting an intricate interplay between plant, microbiota, and nutrient conditions.

Recent studies began to illustrate the importance of immunity-microbiome interplays in plants. For example, we and others have recently shown that PTI-associated PRRs and ROS-generating RBOHD/F are essential components of a plant genetic network in configuring a normal eubiotic leaf microbiota to prevent health-damaging dysbiosis^1,10,11^. Similarly, bacterial members of both leaf and root microbiotas either stimulate or suppress PTI-associated gene expression^50,51^. In this study, we found that a synthetic community composed of 48 culturable *Arabidopsis* phyllosphere bacteria (SynCom^Col-0^) was sufficient to restore immunocompetence in the leaves of AX plants at a level similar to that conferred by a natural soil-derived microbial community. This is interesting considering that most members of the leaf bacterial microbiota live on the surfaces and less than 5% of leaf bacteria reside inside the leaves^10^. Results presented here either suggest the importance of endophytic leaf bacteria in maturing immune responses in *Arabidopsis* leaves or there are multiple functionally redundant sub-communities of any given microbiota and each subcommunity can independently confer immune maturation to plants. In either scenario, there seems to be substantial redundancy among different phyllosphere strains in endowing immunocompetence (Supplementary Fig. 5).

The role of microbiota in modulating immunocompetence seems robust across different plant growth conditions used in our study. When we analyzed the transcriptome profiles of plant colonized by SynCom^Col-0^ vs. natural microbiota (HO^Malaka^ or HO^MSU^), compared to the corresponding axenic plants, enrichment of immune-associated genes was observed in both cases (Supplementary Fig. 8), even though plants were grown under different conditions, including different grown substrate mixtures and photoperiods (see Methods). Interestingly, enriched immune genes observed in our study include 20 so-called “general non-self response (GNSR)” genes that are commonly induced by 13 individual strains from the *At-LSPHERE* collection^52^. The GNSR genes constitute 9 % of the upregulated genes (20/213) commonly enriched in plants inoculated with natural microbiotas and SynCom^Col-0^ in this study. Overall, our study is consistent with the existence of a broader core microbiota-associated transcriptome response and highlights the importance of a natural or eubiotic community in shaping the transcriptome landscape of basal immune responses in plants.

Another important implication of the findings from this study is that not only do *Arabidopsis* plants require a microbiota to properly develop PTI, but also that the composition of the microbiota is important. We found that SynCom^Col-0^, a eubiotic microbiota derived from healthy *Arabidopsis* leaves, was sufficient to restore immunocompetence to AX plants. In contrast, SynCom^mfec^, a dysbiotic microbiota derived from leaves of the *Arabidopsis mfec* quadruple mutant, overstimulated immune gene expression (Fig. 6). This observation suggests that a healthy, eubiotic microbiota is necessary to properly gate the plant immune system. We think that this is an important observation because in human-microbiome interactions dysbiosis is associated with autoimmune ailments such as inflammatory bowel disease, diabetes, allergies and other health issues^53,54^. Thus, an intimate interplay between immunity and microbiota appears to be core to host-microbiome interactions in both animal and plant kingdoms. Deviations from a eubiotic microbiota could result in immunodeficiency (as in the case of AX plants) or immune-overstimulation (as in the case of SynCom^mfec^-inoculated plants). Thus, a eubiotic microbiota has a fundamental role in gating plant immune response during growth and development.

## Methods

### *Arabidopsis* growth conditions

The following *Arabidopsis thaliana* genotypes were used in this study: Col-0, *bak1-5 bkk1-1 cerk1* mutant (*bbc*)^46^. Conventionally grown plants were grown using potting soil composed of equal parts Suremix (Michigan Grower Products), medium vermiculite, and perlite. The resulting potting soil was autoclaved once to eliminate pests. Plants were grown in an air-circulating growth chamber with the following conditions: 60% RH, 22 °C, 12 h day/12 h night photoperiod cycle and provided with a daytime photon flux of ∼90-100 μmol m^−2^ s^−1^ and supplemented with 0.5x Hoagland nutrient solution^55^ as needed.

For peat-based gnotobiotic experiments, plants were grown in FlowPots or GnotoPots. Methods for preparation and inoculation are described in^43^. Nutrients were supplemented with buffered 0.5x LS liquid media (pH 5.7) (Caisson Labs), unless indicated otherwise. Full strength LS contains 1900 mg/L KNO_3_, 1650 mg/L NH_4_NO_3_, 332.2 mg/L CaCl_2_, 200 mg/L MES buffer, 180.7 mg/L MgSO_4_, 170 mg/L KH_2_PO_4_, 100 mg/L myo-inositol, 98 mg/L KHCO_3_, 37.26 mg/L EDTA, 27.8 mg/L FeSO_4_ ⋅ 7H_2_O, 16.9 mg/L MnSO_4_ ⋅ H_2_O, 8.6 mg/L ZnSO_4_ ⋅ 7H_2_O, 6.2 mg/L H_3_BO_3_, 0.83 mg/L KI, 0.4 mg/L thiamine HCl, 0.25 mg/L Na_2_MoO_4_ ⋅ 2H_2_O, 0.025 mg/L CoCl_2_ ⋅ 6H_2_O, and 0.025 mg/L CuSO_4_ ⋅ 5H_2_O. Soil for natural microbiota inoculation was harvested from a Miscanthus plot at Michigan State University (42.716989, −84.462711; ‘MSU’ microbiota input). For the transcriptome experiment using HO communities, a second natural microbiota input was obtained from soil harvested from an undisturbed grassland in Malaka Township, IA (41.836100, −93.007800; ‘Malaka’ microbiota input). For natural community microbiota experiments, AX plants were mock-inoculated with an autoclaved soil slurry (50 g soil/ L water) and HO plants were inoculated with the same unautoclaved soil slurry. For experiments using synthetic communities, plants were inoculated as previously described^10^. Briefly, individual microbiota members were cultured individually on individual R2A (Sigma) plates before being pooled together in equal ratios (OD_600_) in 10 mM MgCl_2_. For GnotoPot assays bacterial suspensions were adjusted to a final OD_600_ = 0.04 (∼2 × 10^7^ cfu/mL) and 1 mL was used to inoculate each GnotoPot. For plate based assays 2 uL of bacterial suspension with final OD_600_ = 0.01 (∼5 × 10^6^ cfu/mL) was spotted directly onto seeds. AX plants for synthetic community experiments were mock-inoculated with an equal volume of 10 mM MgCl_2_.

### Pathogen infection assays

For flg22 protection assays with conventional potting soil-grown *Arabidopsis*, plants of the indicated ages were hand-infiltrated using a blunt-end syringe with 500 nM flg22 and allowed to dry until no longer water-soaked in appearance. 16-24 h after pretreatment with flg22 leaves were infiltrated with 5×10^7^ cfu/mL *Pst* DC3000 using a blunt-end syringe. Infected plants were partially covered with a clear plastic dome to increase humidity. Bacterial populations were determined 24 h after infiltration.

For flg22 protection assays in gnotobiotic *Arabidopsis*, plants were grown in FlowPots with 0.5x LS. Sow date was staggered, and plants of the indicated ages were treated at the same time to allow direct comparison. Plants were pretreated with 100 nM flg22 using a blunt-end syringe and allowed to dry until no longer water-soaked in appearance. Control and flg22-treated plants were kept in microboxes with the lid on overnight. Twenty-four h after flg22 pretreatment, plants were syringe-infiltrated with *Pst* DC3000 at 1×10^6^ cfu/mL. Infected plants were allowed to dry until no longer water-soaked in appearance and then covered with a clear plastic dome to maintain high humidity. Bacterial populations were determined 24 h after infiltration.

For disease assays (without flg22 pretreatment) in gnotobiotic *Arabidopsis*, plants were grown in FlowPots or GnotoPots with 0.5x LS and hand-infiltrated with *Pst* DC3000 at 1×10^5^ cfu/mL. Infected plants were allowed to dry then then kept at high humidity (>95% RH). Bacterial populations were determined three days after infiltration. For *Bc* inoculation, spores were diluted in 1% Sabouraud Maltose Broth to a final concentration of 1 × 10^5^ spores/mL. Two 2-uL droplets were spotted per leaf. Infected plants were kept at high humidity (>95% RH). Lesions were imaged five days after inoculation.

### Transcriptome analysis

For transcriptome experiments with natural community inputs, total RNA was extracted from whole rosettes of FlowPot-grown *Arabidopsis* inoculated with ‘MSU’ ‘Malaka’) soil-derived input microbiota or in the case of AX plants, mock-inoculated with a corresponding input microbiota that had been autoclaved. A biological replicate is defined as a pool of eight rosettes collected from four FlowPots within the same microbox. Three biological replicates per condition were collected, totaling six holoxenic and six axenic replicates. RNA was extracted using the RNeasy Plant Mini Kit (Qiagen) according to manufacturer’s protocol with optional on-column DNase digestion. Purified RNA was eluted in TE buffer (Tris-HCl 10 mM, pH 7.5, EDTA 1 mM). RNA concentrations were determined using an ND-1000 NanoDrop spectrophotometer (Thermo Scientific) or by Qubit RNA HS fluorometric assay (Thermo Fisher). Total RNA samples were collected in 2.0 ml nucleic acid LoBind tubes (Eppendorf) and stored at −80 °C. RNA was checked for quality using a Bioanalyzer 2100 (Agilent), and all samples were determined to have an RNA integrity (RIN) score of six or greater. Stranded sequencing libraries were prepared using the NuGEN Ovation RNA-SEQ System for Model Organisms (*Arabidopsis*) according to manufacturer’s protocol (NuGEN). Library preparation and sequencing was performed by the Michigan State University Research Technology Service Facility (RTSF). Sequencing was performed on the HiSeq 2500 (Illumina) with a 1 x 50 bp single read stranded format using Illumina HiSeq SBS reagents (version 4). Base calling was done by Illumina Real Time Analysis (RTA version 1.18.64).

For transcriptome experiments with SynComs, plants were grown in GnotoPots under long day (16 h day/8h night) condition and sampled at day 26 after germination. RNA extraction was done as described above but samples were eluted in RNase/DNase-free water. RNA quality controls were performed using the Qubit (Thermo Fisher Scientific) and TapeStation (Agilent Technologies, Inc.). Stranded RNA-seq libraries were pooled and sequenced on the Illumina NovaSeq 6000 S1 to obtain 50-bp pair-end reads. Library preparation and sequencing was performed by the Sequencing and Genomic Technologies Core at Duke University’s Center for Genomic and Computational Biology.

Raw transcriptome reads for both transcriptome experiments were processed on the Duke Compute Cluster as follows: Read quality control was performed using FastQC^56^, adapter trimming and sequence mapping was achieved using Trimmomatic^57^ and STAR version 9.3.0^58^. Gene expression was quantified using the R package Rsubreads version 2.8.2^59^. Differentially expressed genes (DEGs) were identified using the R package DESeq2^60^. Read transformation and normalization for PCoA and clustering of was done using the EdgeR package on the iDEP platform (version 1.0)^61^. Genes with differential expression were selected using |log_2_FC| > 1 and FDR < 0.05 (calculated using default DESeq2 settings based on Benjamini–Hochberg corrected Wald test) as selection criteria and GO analysis was performed using ShinyGO version 0.76.2^62^ with an FDR cutoff of 0.05 and 4 gene per group selection criteria.

### ROS burst assay

Leaf disks (4 mm in diameter) were taken from the center of leaves from plants of various ages indicated and floated with abaxial side down in wells of a white 96-well plate containing 200 μL sterile water in each well. Plates were covered with foil and leaf disks were kept in sterile water overnight to attenuate wounding response. Twenty-four h later, water was removed from wells and replaced with 100 μL of an immune-eliciting solution containing 34 μg/mL luminol (Sigma), 20 μg/mL horseradish peroxidase (Sigma), and 100-250 nM of the indicated PAMP/DAMP. Luminescence was measured (total photon counting) over 40 min immediately after the addition of immune-eliciting solution using a SpectraMax L microplate reader (Molecular Devices). Total ROS was calculated in Prism (GraphPad) using ‘Area under curve’ analysis.

### RT-qPCR analysis gene expression

For RT-qPCR analysis of elicitor-induced gene expression, whole plants were sprayed with or leaf disks were floated on an elicitor solution. For spray elicitation (Fig. 1d,e and 2f), plants of the indicated ages were treated with a foliar spray of elicitor solution consisting of 100 nM flg22, 0.1% DMSO, and 0.025% Silwet-L77 (Bioworld) or a mock solution that lacked flg22. Foliar sprays were applied ensuring treatment solution came in contact with both the adaxial and abaxial side of leaves. For leaf disk elicitation (Fig. 3b and Supplementary Fig. 5 b,c), 4 mm leaf disks were taken from 4.5-6-week-old plants and floated on sterile water overnight. The next day the water was removed and replaced with an elicitor solution containing 250 nM of the indicated PAMP/DAMP. For basal gene expression analysis of plate grown plants (Fig. 6a-c), full rosettes of 16-days old seedlings were snipped and transferred to 2 mL screw-top tubes before being frozen in liquid N_2_ and stored at −80 °C until further processing. The above-ground tissue from 5 plants was pooled to constitute one biological replicate. For transcriptional analysis of SynCom leaf infiltration (Supplementary Table 6) 4.5-5 weeks old plants were hand-infiltrated with each strain at OD_600_ 0.2 and three biological replicates were harvested after 24 h for RNA extraction.

Total RNA was extracted from leaf tissues using either Trizol (Thermo Fisher) and a Direct-zol RNA extraction kit (Zymo Research) or an RNeasy Plant Mini Kit (QIAGEN) according to the manufacturer’s instructions using the optional on-column DNase treatment. cDNA synthesis was accomplished in 10 μL volumes with SuperScript IV VILO master mix (Thermo Fisher) or M-MLV Reverse Transcriptase (Thermo Fisher) according to the manufacturer’s instructions using 640-1000 ng total RNA as input. Upon synthesis, cDNA was diluted 10-fold and qPCR was performed in duplicate on a minimum of three biological replicates in 10 μL reaction volumes containing 5 μL SYBR Green PCR master mix, 0.25 μL of each primer, and 2 μL of template cDNA on an ABI 7500 Fast (Applied Biosystems) or QuantStudio 5 (Applied Biosystems) RT-qPCR system using the default settings. *PP2AA3* was used for normalization. The primer sets used to quantify gene expression in this study are listed in Supplementary Table 9.

### Quantification of salicylic acid (SA) and glucosylated salicylic acid (SAG)

Plant hormones SA and SAG were extracted as previously described^63^. In brief, 100 mg of leaf tissue harvested from 4.5-week-old plants grown in FlowPots was frozen and ground to fine powders with a TissueLyser (Qiagen) using two 45 s cycles at 28 Hz. Frozen powders were resuspended in 1 mL extraction buffer containing 80% methanol, 0.1% formic acid, 0.1 mg/mL butylated hydroxytoluene, and 100 nM deuterated abscisic acid (ABA-^2^H_6_) in water. Samples were extracted overnight at 4 °C with gentle agitation. The next day, samples were cleared by centrifugation at 12,000×*g* for 10 minutes, filtered through a 0.2 μm PTFE membrane (Millipore), and transferred to autosampler vials. Ten μL injections of prepared extracts were separated using an Ascentis Express fused-core C18 column (2.1×50 m, 2.7 μm) heated to 50 °C on an Acquity ultra performance liquid chromatography system (Waters Corporation). A gradient of 0.15% formic acid in water (solvent A) and methanol (solvent B) was applied over 2.5 minutes at a flow rate of 0.4 mL/min. Separation consisted of a linear increase from A:B (49:1) to 100% B. Transitions from deprotonated molecules to characteristic product ions were monitored for ABA-^2^H_6_ (*m*/*z* 269.1 > 159.1), SA (*m*/*z* 137.0 > 93.0), and SAG (*m*/*z* 299.1 > 137.0) on a Quattro Premier tandem mass spectrometer (Waters Corporation) in negative ion mode. The capillary voltage, cone voltage, and extractor voltage were 3500 V, 25 V and 5 V, respectively. The flow rates were 50 L/h for the cone gas (N_2_) and 600 L/h for the desolvation gas (N_2_). ABA-^2^H_6_ served as the internal standard for hormone quantification. Collision energies and source cone potentials were optimized using QuanOptimize software. Masslynx v4.1 was used for data acquisition and processing. Peaks were integrated and the analytes quantified based on standard curves normalized to the internal standard.

### Immunoblot analysis

Protein was extracted from leaves as previously described ^19^ with slight modification. First, frozen leaf tissues were ground to fine powders with a TissueLyser (Qiagen) using two 45 s cycles at 28 Hz. Powders were taken up into a protein extraction buffer containing 50 mM Tris-HCl (pH 8.0), 150 mM NaCl, 10% (v/v) glycerol, 1% (v/v) IGEPAL CA-630 (NP-40) (Sigma), 0.5% (w/v) sodium deoxycholate, 1 Complete EDTA-free Protease Inhibitor tablet (Roche) and incubated on ice for 15 min with periodic inversion. Leaf lysates were cleared by centrifugation at 10,000×*g* for 5 min and total protein normalized via Bradford Assay (Biorad). Extracts were prepared for SDS-PAGE with a 5x loading buffer containing 10% (w/v) sodium dodecyl sulfate, 20% glycerol, 0.2 M Tris-HCl (pH 6.8), and 0.05% bromophenol blue and gradually denatured on a thermocycler using the following sequence: 37 °C for 20 min, 50 °C for 15 min, 70 °C for 8 min, and 95 °C for 5 min. Protein was subsequently separated on NuPAGE 4-12% bis-tris gels (Thermo Fisher) for 2.5 h using 100 V. Proteins were then transferred to PVDF using an iBlot 2 dry blotting system (Thermo Fisher), blocked in 3% milk + 2% BSA and immunoblotted overnight at 4 °C with antibodies specific to *Arabidopsis* FLS2 (Agrisera), BAK1 (Agrisera), MPK3 (Sigma) or MPK6 (Sigma) at concentrations recommended by the manufacturer. Blots for detecting phosphorylated MAPK were blocked in 5% BSA and immunoblotted with a phosphor-p44/42 MAPK (Erk1/2) (Thr202/Tyr204) antibody (Cell Signaling). Horseradish peroxidase-conjugated anti-rabbit antibody produced in goat (Agrisera) was used as a secondary antibody and the resulting proteins of interest were visualized with SuperSignal West chemiluminescent substrate (Thermo Fisher). Ponceau S or Amido Black staining was performed to verify equal loading.

### Phyllosphere bacterial enumeration

A culture-based approach was used to quantify phyllosphere bacterial communities as previously described^10^. Briefly, leaves were rinsed in sterile water twice and air dried to remove residual surface water. Leaves were then weighed and ground in 10 mM MgCl_2_ and a serial dilution was plated on R2A containing 50 ug/mL cycloheximide. Plates were incubated at room temperature for two days, then 4 °C for four days and colonies counted.

### Microbial community profiling

16S rRNA gene amplicon sequencing was used to estimate relative abundance of bacterial taxa. Total DNA was extracted from phyllosphere and input communities using DNeasy PowerSoil Pro Kit (Qiagen) according to the manufacturer’s instructions. For phyllosphere samples, 2-3 leaves were pooled from a single plant per biological sample (*n* = 12). For input samples, 500 uL of soil slurry was saved during inoculation (*n* = 5). PCR was performed with AccuPrime high-fidelity Taq DNA polymerase using barcoded primers with heterogeneity adapters targeting the v5/v6 region of the 16S rRNA gene (799F and 1193R, see Supplementary Table S9 for primer sequences). Primary amplicons were separated via electrophoresis on a 1% agarose gel. DNA in the ∼400 bp band was recovered using the Zymoclean Gel DNA Recovery Kit (Zymo Research). The concentration of the recovered DNA was measured with PicoGreen dsDNA assay kit (Invitrogen) and normalized to 1 to 10 ng ul^−1^. Samples were submitted to the Research Technology Service Facility (RTSF) Genomics Core at Michigan State University for library preparation and 16S rRNA gene sequencing.

The RTSF Genomics Core performed secondary PCR using dual indexed, Illumina compatible primers which target the Fluidigm CS1/CS2 oligomers at the ends of your primary PCR products. Amplicons were batch normalized using Invitrogen SequalPrep DNA Normalization plates and recovered product was. The pools were QC’d and quantified using a combination of Qubit dsDNA HS, Agilent 4200 TapeStation HS DNA1000 and Invitrogen Collibri Library Quantification qPCR assays.The library pool was loaded onto a MiSeq v2 flow cell and sequencing performed in a 2×250bp paired end format using a MiSeq v2 500 cycle reagent cartridge. Custom sequencing and index primers complementary to the Fluidigm CS1 and CS2 oligomers were added to appropriate wells of the reagent cartridge. Base calling was done by Illumina Real Time Analysis (RTA) v1.18.54 and output of RTA was demultiplexed and converted to FastQ format with Illumina Bcl2fastq v2.20.0.

Raw fastq files from the MiSeq instrument were demultiplexed and processed using the QIIME 2 Core 2022.2 distribution^64^. In brief, primers and heterogeneity spacers were removed using Cutadapt^65^ and DADA2^66^ was used to trim, quality filter and denoise sequences, remove chimeric sequences and obtain ASVs. Taxonomic assignment of each ASV was performed using a Naïve Bayes classifier^67^ pre-trained on the SILVA 16S rRNA gene reference database (release 138)^68^ formatted for QIIME using RESCRIPt^69^. Unassigned sequences or sequences identified as plant chloroplast or mitochondria were removed. Diversity analyses were performed within QIIME 2. Samples were rarified to 5765 reads for calculating diversity metrics.

### *CYP71A12pro:GUS* histochemical assay

GUS assay was performed as described previously^70^ with minor modifications. Briefly, seedlings were grown in 24-well plates containing liquid LS medium supplemented with 0.5 % sucrose under 16 h/8 h day/night cycle in a Percival chamber at 22 °C under light intensity of 50 μmol m^−2^ s^−1^. Plants were inoculated at day 12 with bacterial strains. Bacterial strains were grown on R2A plates at 22 °C for 3 days, resuspended in 10 mM MgCl_2_, and added to seedlings in LS medium without sucrose at OD_600_ of 0.002. After treatment with SynCom strains for 5 hours, seedlings were rinsed with 0.5 mL of 50 mM sodium phosphate buffer (pH 7) and submerged in 0.5 mL GUS staining solution (50 mM sodium phosphate pH 7, 0.5 mM K_4_[Fe(CN)_6_], 0.5 mM K_3_[Fe(CN)_6_], 1 mM X-Gluc (GoldBio), and 0.01% Silwet L-77). After vacuum-infiltration for 10 min, plates were incubated at 37 °C overnight. Plants were fixed with a 3:1 ethanol:acetic acid solution at 4 °C for 1 day followed by transfer to 95% ethanol.

## Data availability

The RNA-seq raw sequencing and analyzed data have been deposited in the NCBI Gene Expression Omnibus database under accession GSE218961 and GSE218962. Raw source 16S rRNA gene sequences from this project are available in the Sequence Read Archive database under BioProject PRJNA977816, accession numbers SAMN35534885 to SAMN35534914. Source data are provided with this paper.

## Code availability

The code used for RNA-seq raw data analysis can be found at: https://github.com/rsohrabi/MIP_ms. The entire sequence analysis workflow for 16S amplicon analysis is available at: https://github.com/BradCP/A-critical-role-of-a-eubiotic-microbiota-in-gating-proper-immunocompetence-in-Arabidopsis.

## Acknowledgements

We thank undergraduates Caleigh Griffin, Trevor Ulrich, Franchesca Dion, and Timothy Johnson and lab technician David Rhodes for assistance in gnotobiotic system design and construction and for performing critical experiments that led to these published results. We thank Dr. Hailing Jin for sharing *B. cinerea*. We thank Dr. Fredrick Ausubel for sharing the *CYP71A12_Pro_:GUS* line seeds. We thank members of the He Lab for critical reading of the manuscript. S.Y.H. is an Investigator at Howard Hughes Medical Institute.

## Author information

These authors contributed equally: Bradley C. Paasch, Reza Sohrabi, James M. Kremer.

### Authors and Affiliations

*Department of Biology, Duke University, Durham, NC, USA*

Bradley C. Paasch, Reza Sohrabi, Kinya Nomura & Sheng Yang He

*Howard Hughes Medical Institute, Duke University, Durham, NC, USA*

Bradley C. Paasch, Reza Sohrabi, Kinya Nomura & Sheng Yang He

*Department of Energy Plant Research Laboratory, Michigan State University, East Lansing, MI, USA*

James M. Kremer & Jennifer Martz

*Department of Plant Pathology, University of Georgia, Athens, GA, USA*

Brian Kvitko

### Contributions

J.M.K., J.M.T., and S.Y.H. conceptualized the initial project.

B.C.P., R.S., J.M.K., J.M.T., and S.Y.H. designed the study and analyzed the data.

B.C.P. performed gnotobiotic plant assays and 16S analyses.

R.S. performed gnotobiotic plant assays and RNA-seq analyses.

J.M.K. performed RNA-seq in FlowPots and preliminary gnotobiotic plant assays.

K.N. performed temporal assays in conventional and gnotobiotic plants.

Y.T.C. provided 16S PCR primers with heterogeneity spacers and generated primary amplicons.

J.M. assisted nutrient gnotobiotic assays.

B.K. performed temporal flg22 protection assays in conventionally grown plants. B.C.P., R.S., and S.Y.H. wrote the manuscript with input from all authors.

