## Supplementary Figures for "A critical role of a eubiotic microbiota in gating proper immunocompetence in *Arabidopsis*"

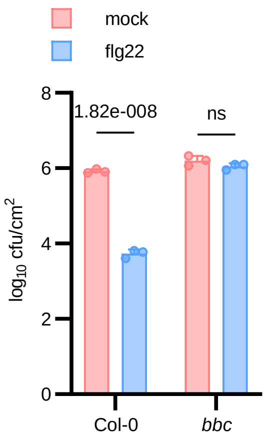


**Supplementary Figure 1 | Holoxenic *bbc* plants do not show robust flg22 protection.**

4-week-old wildtype (Col-0) and *bbc* mutant HO plants were treated 24 hours prior to inoculation with *Pst* DC3000 ($OD_{600}$ = 0.002) with either a water (mock) or 100 nM flg22 solution. Each column represents bacterial titer 24 hours after inoculation as log transformed $cfu/cm^2$ and is the mean of three plants. Error bars indicate SD. (Col-0: $p = 1.82 \times 10^{-8}$, bbc: not significant; two-way ANOVA with Tukey's HSD post-hoc test). This experiment was repeated three times with similar results.

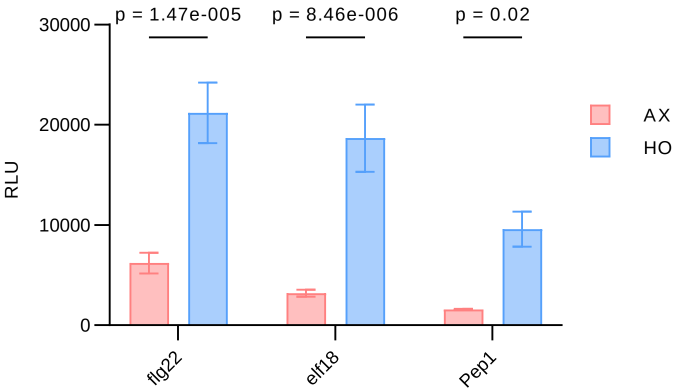


**Supplementary Figure 2 | Axenic *Arabidopsis* plants exhibit decreased total ROS production upon PTI elicitation compared to holoxenic plants.**

Total ROS production induced by 250 nM flg22, elf18, and Pep1 in AX and HO plants in GnotoPots. Results calculated from data presented in Fig. 2a by determining the mean area under curve of the eight plants ± SEM (flg22: $p = 1.47 \times 10^{-5}$, elf18: $p = 8.46 \times 10^{-6}$, Pep1: $p = 0.02$; two-way ANOVA with Šidák multiple comparison test). This experiment was repeated three times with similar results.

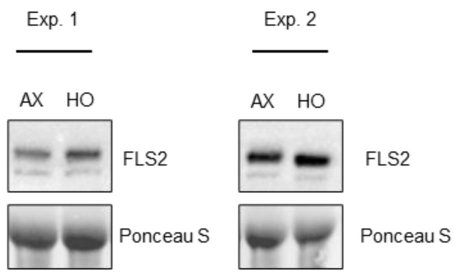


**Supplementary Figure 3 | FLS2 protein abundance in axenic and holoxenic plants.**

Total FLS2 protein detected in whole leaf tissue lysate of four pooled plants. Two experimental repeats show variability in FLS2 relative abundance. Ponceau S stain of all blots show equal loading. This experiment was repeated five times with variable results. Blots from two representative experiments shown. See Source Data Figures 4 and 5 for image cropping.

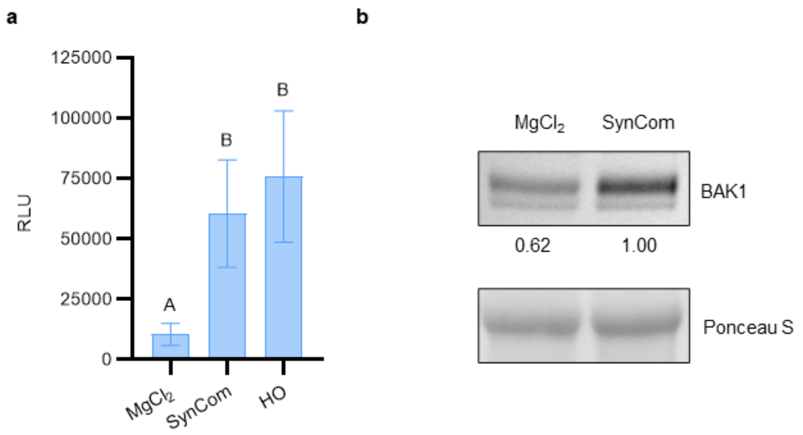


**Supplementary Figure 4 | SynCom$^{Col-0}$ restores immunocompetence.**

**a**, Total ROS production induced by 100 nM flg22 in plants colonized by HO or SynCom$^{Col-0}$. Results calculated from data presented in Fig. 3a by determining the mean area under curve of the 12 plants ± SEM. Different letters represent a significant difference ($p < 0.05$, two-way ANOVA with Tukey's HSD post-hoc test). **b**, Total BAK1 protein detected in leaf lysates of 6-week-old plants mock-inoculated with 10 mM $MgCl_2$ and plants colonized by SynCom$^{Col-0}$. Numbers below blot indicates band intensity relative to that of Ponceau S, normalized to HO = 1.00. These experiment were repeated two times with similar results. See Source Data Figure 6 for image cropping.

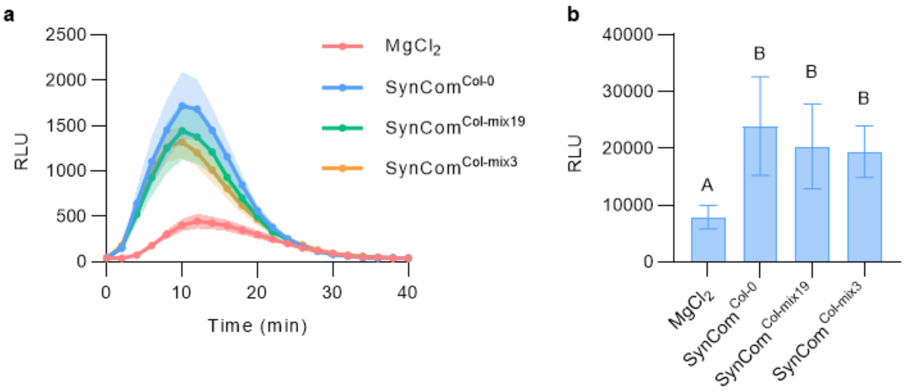


**Supplementary Figure 5 | A simplified SynCom restores immunocompetency.**

**a,** ROS burst kinetics and **b,** total ROS production after induction by 250 nM flg22 in plants colonized by SynCom$^{Col\text{-}mix19}$, SynCom$^{Col\text{-}mix3}$ or mock-inoculated with 10 mM $MgCl_2$ as a control in GnotoPots. Error bars indicate SD. Different letters represent a significant difference ($p < 0.05$, one-way ANOVA with Tukey's HSD post-hoc test). This experiment was repeated two independent times with similar results.

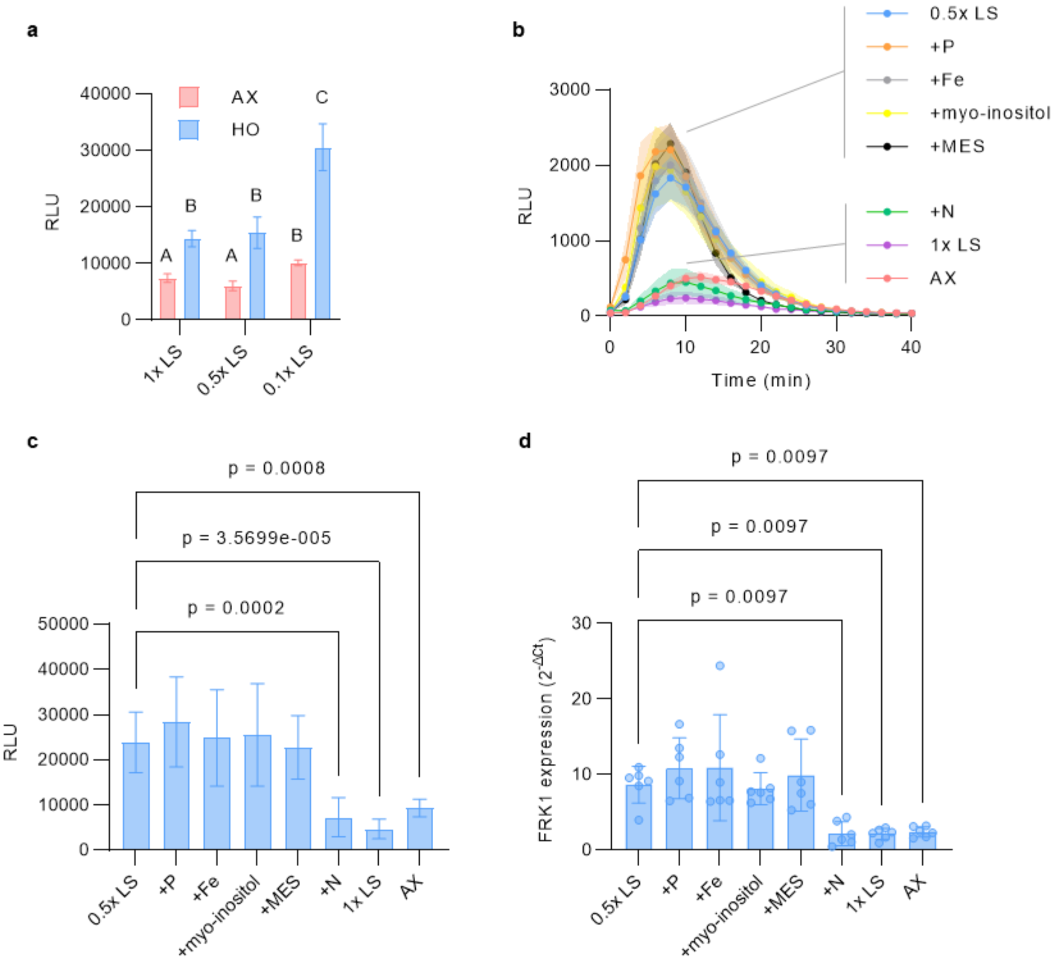


**Supplementary Figure 6 | Excess nutrients suppress microbiota-mediated immune maturation.**

**a**, Total ROS production induced by 100 nM flg22 in AX and HO plants grown in GnotoPots supplied with 0.1x, 0.5x, or 1x LS nutrient solution concentrations. Results calculated from data presented in Fig. 4e by determining the mean area under curve of the six plants ± SEM. Different letters represent a significant difference ($p < 0.05$, Fisher's Least Significant Difference Test). This experiment was repeated three times with similar results. **b-c**, ROS burst dynamics (b) and total ROS (c) induced by 250 nM flg22 in HO plants grown in GnotoPots supplied with 0.5x LS, 0.5x LS supplemented with additional components of LS up to 1x, and 1x LS. AX plants included as a control. Results represent the mean of eight plants ± SEM. Total ROS production calculated by determining the mean area under curve ± SEM (compared to 0.5x LS: $q = 0.0002$ (+N), $q = 3.57 \times 10^{-5}$ (1x LS), $q = 0.0008$ (AX), all others ns; one-way ANOVA with Benjamini-Hochberg FDR correction). **d**, *FRK1* gene expression in AX and HO plants induced by 250 nM flg22. Total RNA was extracted from leaf disks 1.5 h after treatment. *PP2AA3* was used for normalization. Bars represent the mean value of eight plants ± SD (compared to 0.5x LS: $q = 0.0097$ (+N), $q = 0.0097$ (1x LS), $q = 0.0097$ (AX), all others ns; one-way ANOVA with Benjamini-Hochberg FDR correction).

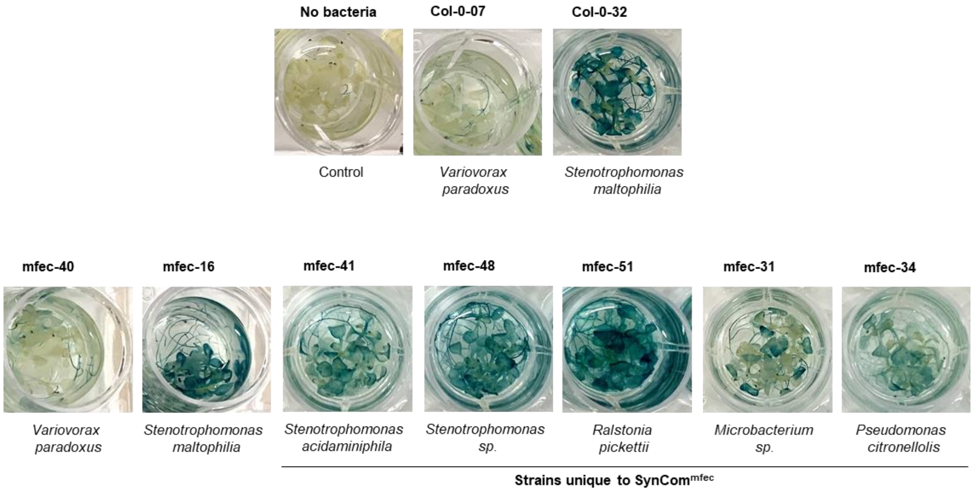


**Supplementary Figure 7 | Induction of *CYP17A12* gene expression by individual members of SynCom$^{\text{Col-0}}$ and SynCom$^{\text{mfec}}$ *CYP71A12*$_{pro}$*:GUS* reporter line.**

GUS histochemical staining was performed after treatment of 12-days old seedling of *CYP71A12*$_{pro}$*:GUS* reporter line with individual SynCom strains. Representative pictures of plants after GUS assay are depicted here. This experiment was repeated two independent times with similar results.

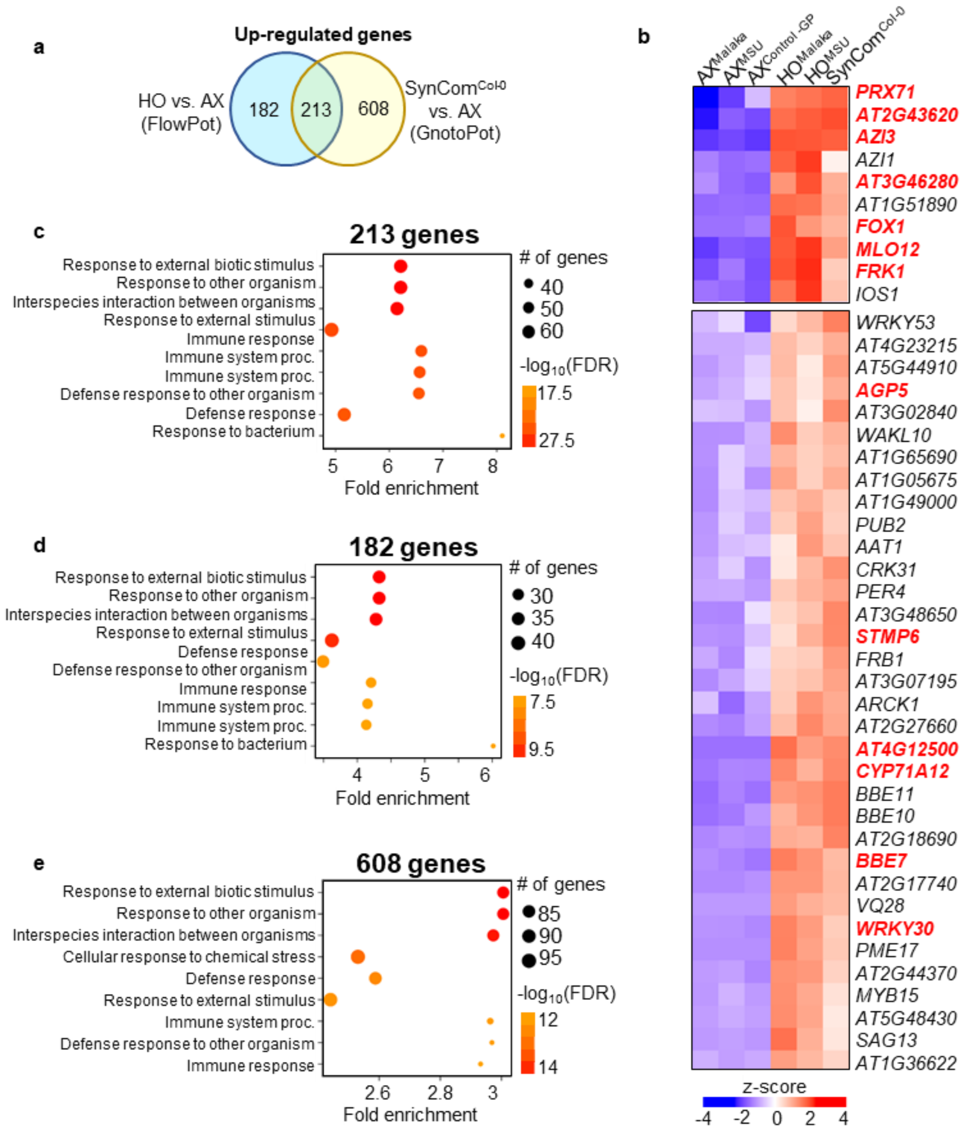


**Supplementary Figure 8 | Leaf transcriptomes of plants colonized with natural community and SynCom$^{Col-0}$ share common immune-related gene expression.**

**a.** Venn diagram of upregulated DEGs showed 213 common Arabidopsis genes in response to natural microbiota and SynCom$^{Col-0}$ colonization. Significant DEGs were identified using DESEq2 with $|\log_2 FC| > 1$ and FDR < 0.05 (Benjamini-Hochberg corrected Wald Test) criteria in a comparison of HO plants (colonized by microbial communities from two different locations/soil types 'MSU' and 'Malaka') and SynCom$^{Col-0}$-colonizedplants with their corresponding AX control. **b**, A subset of the differentially regulated genes in HO and SynCom$^{\text{Col-0}}$ plants, compared to corresponding AX plants, is shown. Heat map of the DEGs was generated using hierarchical clustering with Euclidean distance and complete linkage. **c-e,** Gene Ontology (GO) term enrichment (GO:BP biological process) analysis on 213 common enriched DEGs in both HO and SynCom$^{\text{Col-0}}$, only in HO or only in SynCom$^{\text{Col-0}}$ plants, compared to their respective AX control plants. Top enriched GO terms are displayed, ranked by significance (FDR adjusted hypergeometric test). The 213 common enriched DEGs in both HO and SynCom$^{\text{Col-0}}$ showed highest fold enrichment (>5) for immunity associated GO terms (Supplementary Fig. 8c). GNSR genes present in 213 common DEGs in both HO and SynCom$^{\text{Col-0}}$, are marked in red.
